# Biomechanical influence of plate configurations on mandible subcondylar fracture fixation: a finite element study

**DOI:** 10.1101/2022.12.12.520039

**Authors:** Anoushka Gupta, Abir Dutta, Kaushik Dutta, Kaushik Mukherjee

## Abstract

Mandible subcondylar fractures have very high complication rate, yet there is no consensus in a suitable plate design for optimal patient outcomes. The present study is aimed at understanding the subcondylar fracture fixation by comparing load transfer in intact and reconstructed fractured mandibles with five different plates: single mini, trapezoid, lambda, strut, and double-mini plates under the complete mastication cycle. Under contralateral molar occlusion (LMOL), the single mini plate resulted in the highest strains. On the contrary, during ipsilateral molar clenching (RMOL), the tensile and compressive strain distributions were found to be reversed, with the tensile strains at the posterior border resulting in lesser strain in reconstructed mandible with single mini plate. Owing to the reduced strains in the reconstructed mandibles, the contralateral molar clenching task is preferred during the immediate post-surgery period for patients. Under this contralateral molar clenching, the peak von Mises stresses in the plate decreased with increase in the number of screws. Furthermore, the presence of two arms seems beneficial to neutralise the tensile and compressive strains across load cases. Consequently, double mini and trapezoid plates were found to perform better as compared to single mini plate during the entire mastication cycle for subcondylar fracture fixation.

## 1. Introduction

Mandible fractures are the most frequent facial skeleton fractures requiring surgery^1^, with the most common type being the subcondylar fractures^2^. Majority of these fractures are caused by traffic accidents, violent assaults, falls, and sports injuries leading to immediate surgical measures^3^. In recent years, the open reduction and internal fixation (ORIF) has gained reputation as the preferred treatment strategy for restoration of anatomical alignment in mandible fractures^3^. However, this fixation method also leads to post-operative complications and expensive revision surgeries in 10-30% of the patients^4^. Surgeons opting for the ORIF of subcondylar fractures, face challenges while identifying a suitable plate configuration for the reconstruction^5^. The geometrical narrow shape of the subcondylar neck and presence of delicate facial nerve near fracture site pose restrictions on the usage of multiple plates for achieving a stable fixation^6^.

Several studies have reported that a single mini plate, placed along condylar axis, provided insufficient structural stability to the fracture fixation^7–9^, resulting in high complication rate of up to 33%^10, 11^. In 2007, Meyer et al. ^12^ defined the ideal osteosynthesis lines along the anterior tensile strains based on experimental models. It increased the usage of double mini plates with an additional plate placed just below the mandibular notch along the ideal osteosynthesis lines^12^. Recently, three new types of three-dimensional plates, lambda, strut and trapezoid plates, were designed to neutralise both the anterior and posterior strain contours and minimize the exposure of the underlying fractured area required for the surgery in the narrow subcondylar area^13^.

Although there are multiple finite element (FE) studies^14–17^ investigating the efficacies of these plating techniques, most of such studies included simulation of only single bite force^14, 16, 17^ or selected loading cases from the mastication cycle^5, 15^ while neglecting the ‘alternating bilateral clenching cycle’^18^. Another common limitation is ignoring the periodontal ligament (PDL) and teeth^14, 19, 20^. The present study is aimed at understanding the subcondylar fracture fixation by comparing load transfer in intact and reconstructed fractured mandibles with five different plates: trapezoidal, lambda, strut, single and double-mini plates whole considering the complex mandible structure and the mastication cycle. Furthermore, the influence of plate designs on important biomechanical parameters such as stresses and interfragmentary displacement was assessed.

## 2. Materials and Methods

Anonymized computed tomography (CT) data of a 20-year-old male, with 512 × 512 pixels resolution, 0.439 mm pixel size and 0.75 mm slice thickness, were used to develop the intact mandible model. Five types of fixation strategies for typical subcondylar fracture were simulated. For a systematic comparison, all the models were modelled with the same methodology as described below.

### 2.1 Development of FE model of mandible

The 3D model (Figure 1) of an intact human mandible was reconstructed using the image processing software Mimics 24.0 (Materialise, Leuven, Belgium). The model had five components: teeth, periodontal ligament (PDL), fibrocartilage, cancellous and cortical bone. Despite the segmentation of the ~0.2 mm thin PDL being challenging ^21^, the PDL was included in the present study since it had notable impact on the stress and strain distributions in mandible^22, 23^. The mandibular condyle articulated against a 0.3 mm fibrocartilage layer attached to the glenoid fossa in solid blocks (20 × 15 × 15 mm), representing the temporal bone^22, 24^. The model was meshed with quadratic tetrahedral element in ANSYS ICEM CFD software (ANSYS Inc., Canonsburg, PA)^25^. A mesh convergence study was performed on intact mandible models comprising of 10,75,810, 13,71,606 and 14,88,538 numbers of elements. The difference of maximum tensile strain was 3% between 1^st^ and 2^nd^ models. However, the difference increased to 6% between 2^nd^ and 3^rd^ models. Hence, the 2^nd^ model, with an average element edge length of 0.2 to 1 mm, was chosen for further analysis.

**Figure 1.**
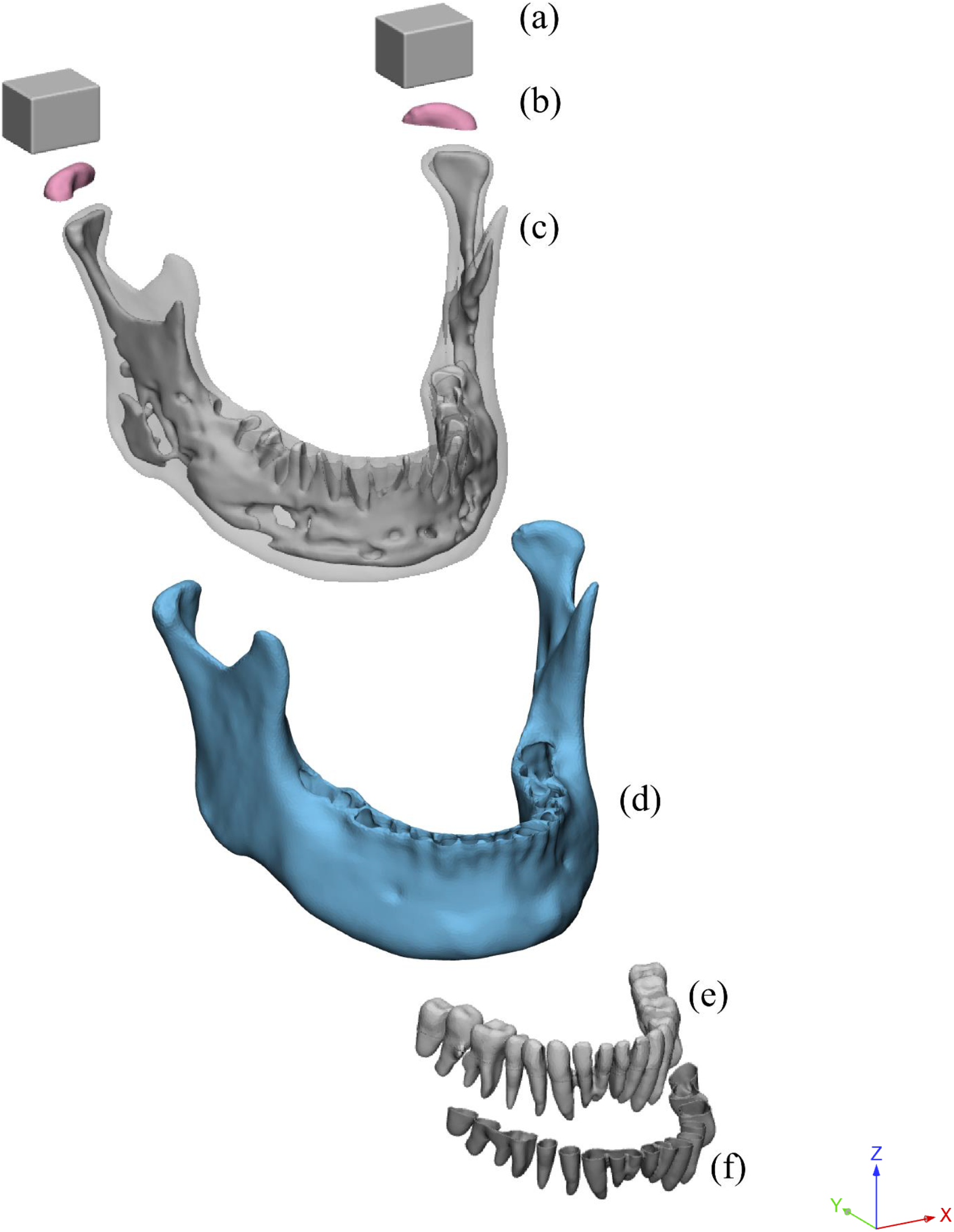
Different components of the mandible model: (a) Blocks, (b) Fibrocartilage on the articular condyle, (c) Cancellous bone (opaque shown inside transparent cortical bone), (d) Cortical bone (opaque), (e) Teeth, and (f) Periodontal ligament (PDL).

### 2.2 Development of implanted mandibles

The unilateral right subcondylar fracture was simulated by following the standard osteotomy line^12^ from the bottom of the sigmoid notch to the midpoint of the posterior ramus on the right side of the mandible. The following types of plates were adapted based on the commercially available plates (DePuy Synthes, PA, United States): 4-hole trapezoid plate, 7-hole lambda plate, 5-hole strut plate and the conventional 4-hole single and double mini plates. The plates were modelled in 3-Matic v12.0 (Materialise, Leuven, Belgium) with a thickness of 1 mm (Figure 2). Screws were modelled as cylinders with 2 mm diameter and 6 mm length. The single mini plate was placed parallel to the condylar axis at the posterior border^13^. The double mini plates were implanted in an angulated manner, i.e., the second mini plate was placed along anterior tensile line^12^. The trapezoidal, lambda and strut plates were placed based on the clinical recommendations^13^. These implanted models were meshed using ~14,00,000 quadratic tetrahedral elements.

**Figure 2.**
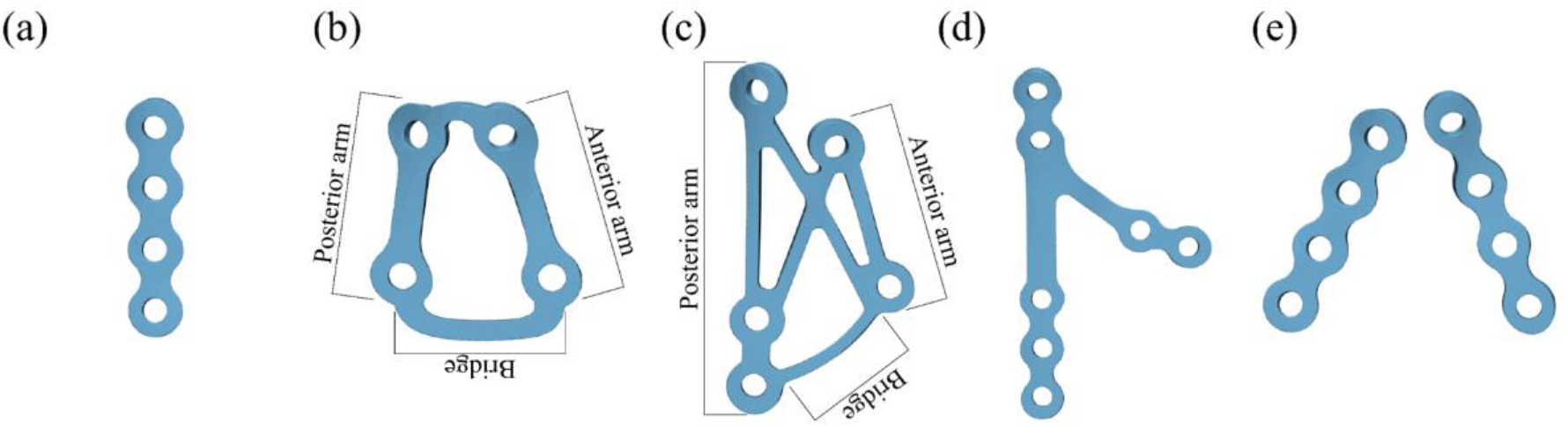
Developed models of different plating techniques: (a) Single mini, (b) Trapezoid, (c) Strut, (d) Lambda, (e) Double mini plates

### 2.3 Material Properties

Region-specific linear elastic orthotropic material properties were assigned to the cortical bone ^24^, and all the other components were modelled as linear elastic isotropic material as given in Table 1^21, 24^. The heterogeneity in the cancellous bone was captured through assigning voxel-based apparent density and Young’s modulus^21^. All the negative grey values were assigned with the minimum elastic modulus^26–28^. A power-law relationship^29^ was used to calculate the Young’s modulus (E) of the cancellous bone: E = 2017.3 × ρ^2.46^. The Poisson’s ratio for cancellous bone was assigned as 0.3^30^. The blocks, representing the temporal bone, were assumed to be rigid. The plates and screws were assumed to be made of titanium alloy (Ti-6Al-4V; E = 110 GPa and ν = 0.3)^21^.

**Table 1.** Material properties of every part of the finite element model

| *Mandibular components* | *Young's Modulus (GPa)* |  |  | *Poisson's Ratio* |  |  | *Shear Modulus (GPa)* |  |  |
| --- | --- | --- | --- | --- | --- | --- | --- | --- | --- |
| | $E_X$ | $E_Y$ | $E_Z$ | $\nu_{XY}$ | $\nu_{YZ}$ | $\nu_{XZ}$ | $G_{XY}$ | $G_{YZ}$ | $G_{XZ}$ |
| Symphyseal and mental | 23 | 10 | 15 | 0.3 | 0.3 | 0.3 | 4.8 | 3.6 | 6.2 |
| Gonial angle | 20 | 11 | 12 | 0.3 | 0.3 | 0.3 | 4.8 | 5.3 | 6.0 |
| Rest of cortical | 17 | 6.9 | 8.2 | 0.3 | 0.3 | 0.3 | 2.8 | 2.9 | 4.6 |
| Cancellous | $10^{-4} - 4.3$ | - | - | 0.3 | - | - | - | - | - |
| Enamel | 80 | - | - | 0.3 | - | - | - | - | - |
| Dentin | 17.6 | - | - | 0.25 | - | - | - | - | - |
| Condylar Fibrous Tissue | 0.006 | - | - | 0.25 | - | - | - | - | - |
| Periodontal ligament | 0.003 | - | - | 0.45 | - | - | - | - | - |
| Plates and Screws (Ti-6Al-4V) | 110 | - | - | 0.3 | - | - | - | - | - |

### 2.4 Applied loading and boundary conditions

Six load cases representing a complete bilateral clenching cycle^31, 32^ were considered here. These load cases represented right molar clenching (RMOL; ipsilateral to the fracture), left molar clenching (LMOL; contralateral to the fracture), left group function (LGF), right group function (RGF), incisal clench (INC) and intercuspal clenching (ICP)^18^. The activity of the following seven muscles were considered: superficial and deep Masseter (SM and DM); lateral and medial Pterygoid (LP and MP); and anterior, middle, and posterior Temporalis (AT, MT and PT). The magnitude and direction of forces were based on the available literature^18^. The individual muscle forces were evenly distributed on nodes representing the corresponding muscle attachment sites. The top of the blocks, representing the temporal bone were constrained in all directions during the mastication cycle. The details of the prescribed loading condition are presented elsewhere^18, 21^. The fracture and bone-plate interfaces were modelled as frictional contact pair with a frictional coefficient of 0.4^33^. An augmented Lagrange algorithm was used to solve the contact problem. The screw surfaces were modelled as ‘bonded’ with the plate, cortical and cancellous bone. Interfaces between various mandible components were modelled as bonded.

### 2.5 Verification and Interpretation of the results

Direct one-to-one experimental validation of the FE model was not possible since the used CT scan dataset was from a living subject. However, a qualitative and quantitative comparison of results were performed for the verification of the models. The occlusal force generated during right molar clench for healthy mandible was 851.6 N, which was within the range of previously reported bite forces^34^. The overall deformation of the mandible was found to be towards the side where the teeth were constrained, similar to those reported earlier^21, 24^.

Analysing the state of strains is important as based on the mechanostat theory, strains influence healing of the fracture site ^35^. In all the load cases, high tensile and compressive strains (~700-940 με) were observed. These strain levels were comparable with those reported in earlier studies ^21, 24^ having similar boundary and loading conditions. These corroborations provided confidence in the developed FE models.

The principal stresses were calculated to predict the failure of cortical bone. The maximum von Mises stresses were calculated for the implants and compared with yield limit of titanium alloy (Ti-6Al-4V) to predict the implant failure. The 95^th^ percentile values of the stresses and strains were calculated to avoid the artificial stress and strain concentrations. Moreover, the interfragmentary displacement at the fracture site was calculated based on maximum opening at the anterior end (Figure 3).

**Figure 3.**
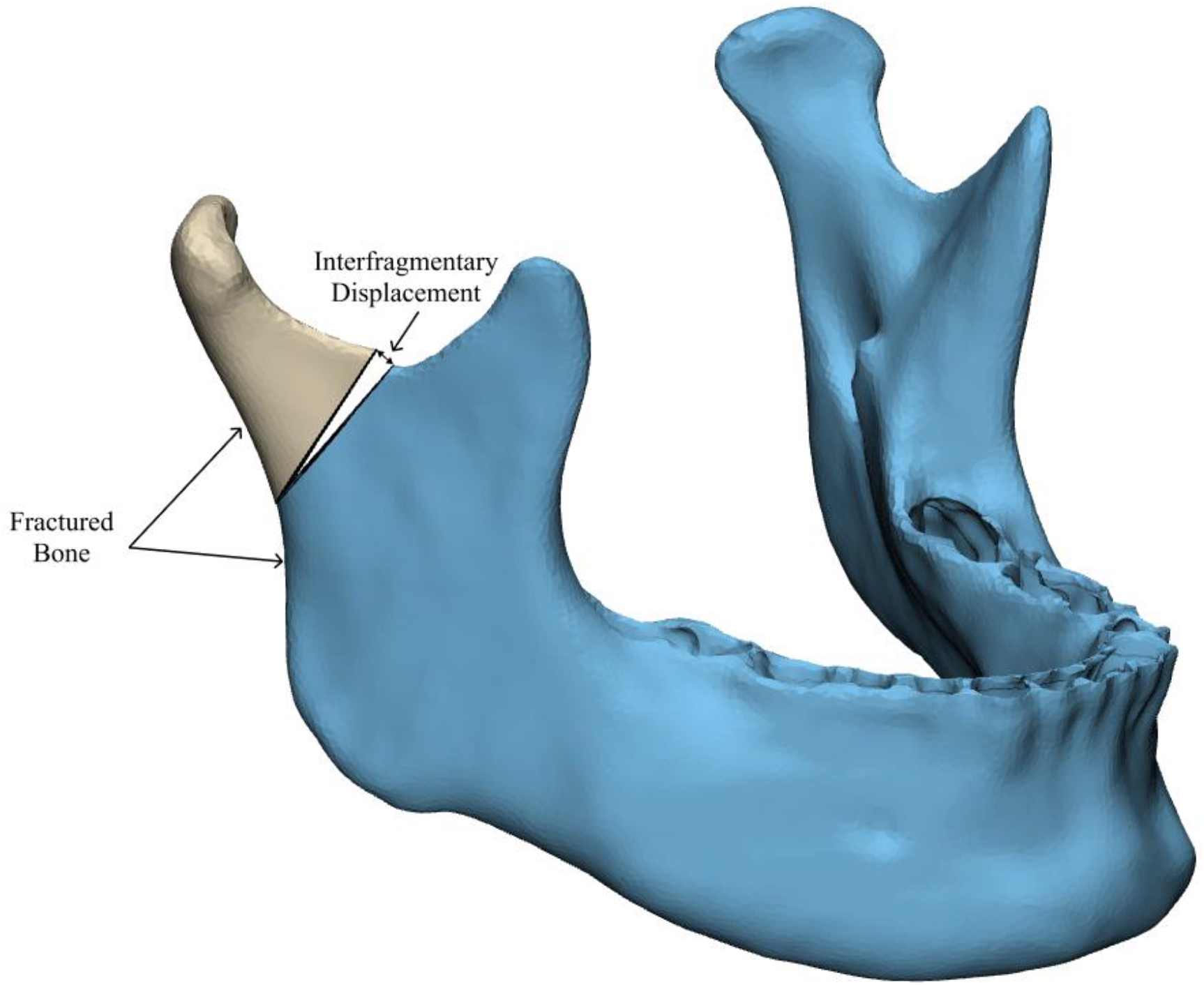
The interfragmentary displacement as the maximum gap at the anterior end at the osteotomy site

## 3. Results

The principal stresses and strains in the mandible, and von Mises stresses in the implants (plates and screws) along with the interfragmentary displacements during molar clenching are discussed below. Findings during the molar clenching tasks are presented for brevity, whereas results corresponding to the rest of the mastication cycle are available in supplementary information.

### 3.1. Stresses and strains in Mandibles

The maximum principal tensile stresses in the intact mandible were observed as ~ 11 – 15 MPa during the mastication cycle. The high maximum principal tensile stress (~14 – 15 MPa) in the intact mandible was found around the molar teeth on the buccal border of mandibles during the molar clenching tasks, with a higher value for RMOL as compared to LMOL. The tensile stress regions were more extensive in the reconstructed mandibles (~17 – 18 MPa) than the intact mandible with the contours running continuously from the fractured region towards the bottommost border of the cortical bone under RMOL (Figure 4). Whereas the high tensile stresses (~15 – 16 MPa) were found near the coronoid of the reconstructed mandibles away from the fracture side under LMOL (Figure 4h-4l), and the least stresses (~15 MPa) were found with lambda and strut plate (Figure 4i-j). Maximum von Mises stresses in the cancellous were found to be ~4 – 8 MPa during the mastication cycle.

**Figure 4.**
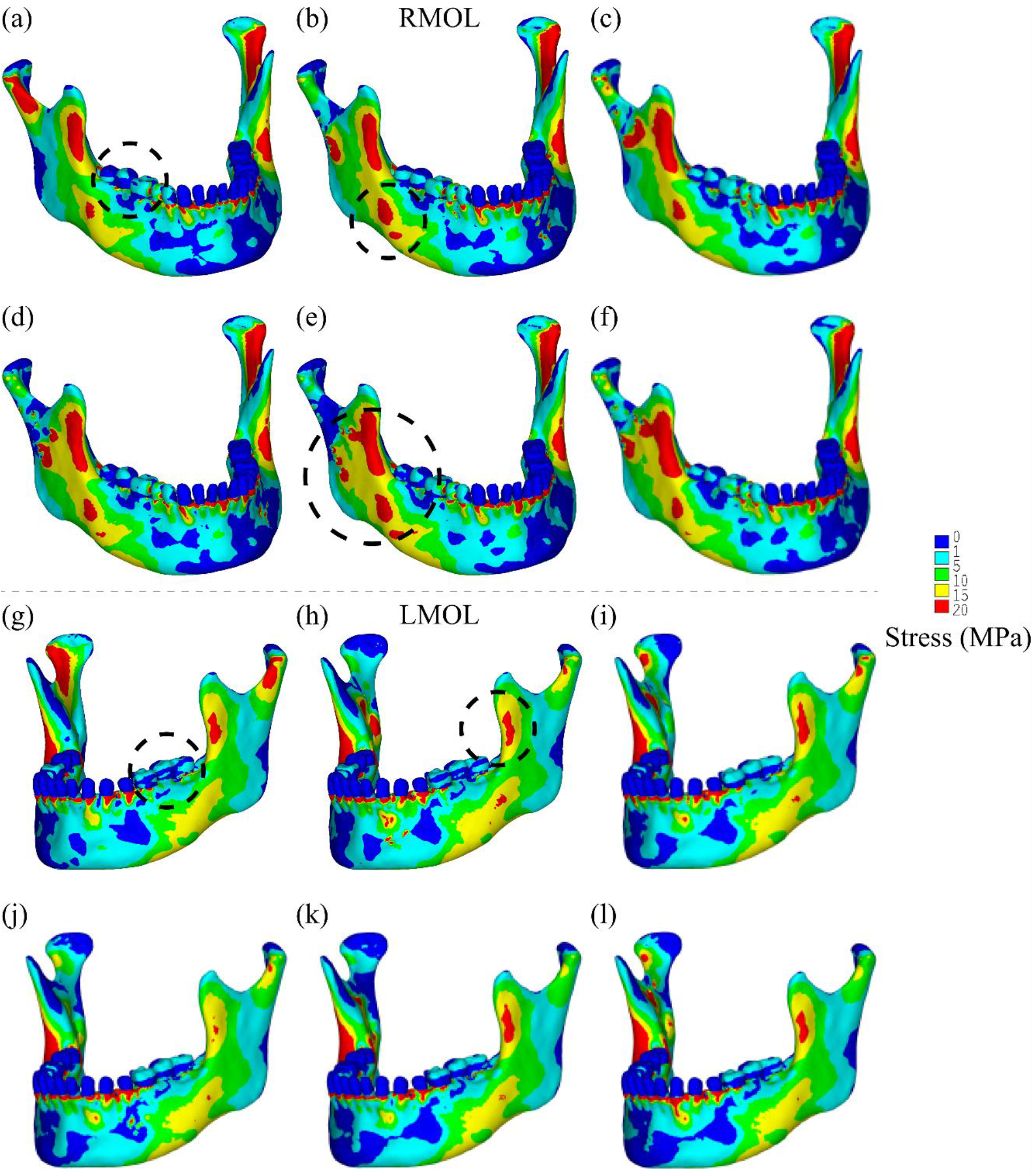
Principal tensile stress distributions: (a, g) intact and reconstructed mandibles with, (b, h) single mini, (c, i) trapezoid, (d, j) strut, (e, k) lambda, (f, l) double mini plates under RMOL and LMOL

The maximum principal tensile and compressive strains in the intact mandible during the mastication cycle were found to be ~700 – 980 με. The molar clenching tasks (RMOL and LMOL) induced high tensile strains of ~860 – 940 με around the condyle and coronoid, towards the lower buccal sides of the intact mandible (Figure 5 a, g). The molar clenching tasks resulted in the highest maximum compressive strain (~870 με) located near the molar teeth. High compressive strains were also found at the anterior border towards the condylar and the coronoid areas (Figure S1 and S2).

**Figure 5.**
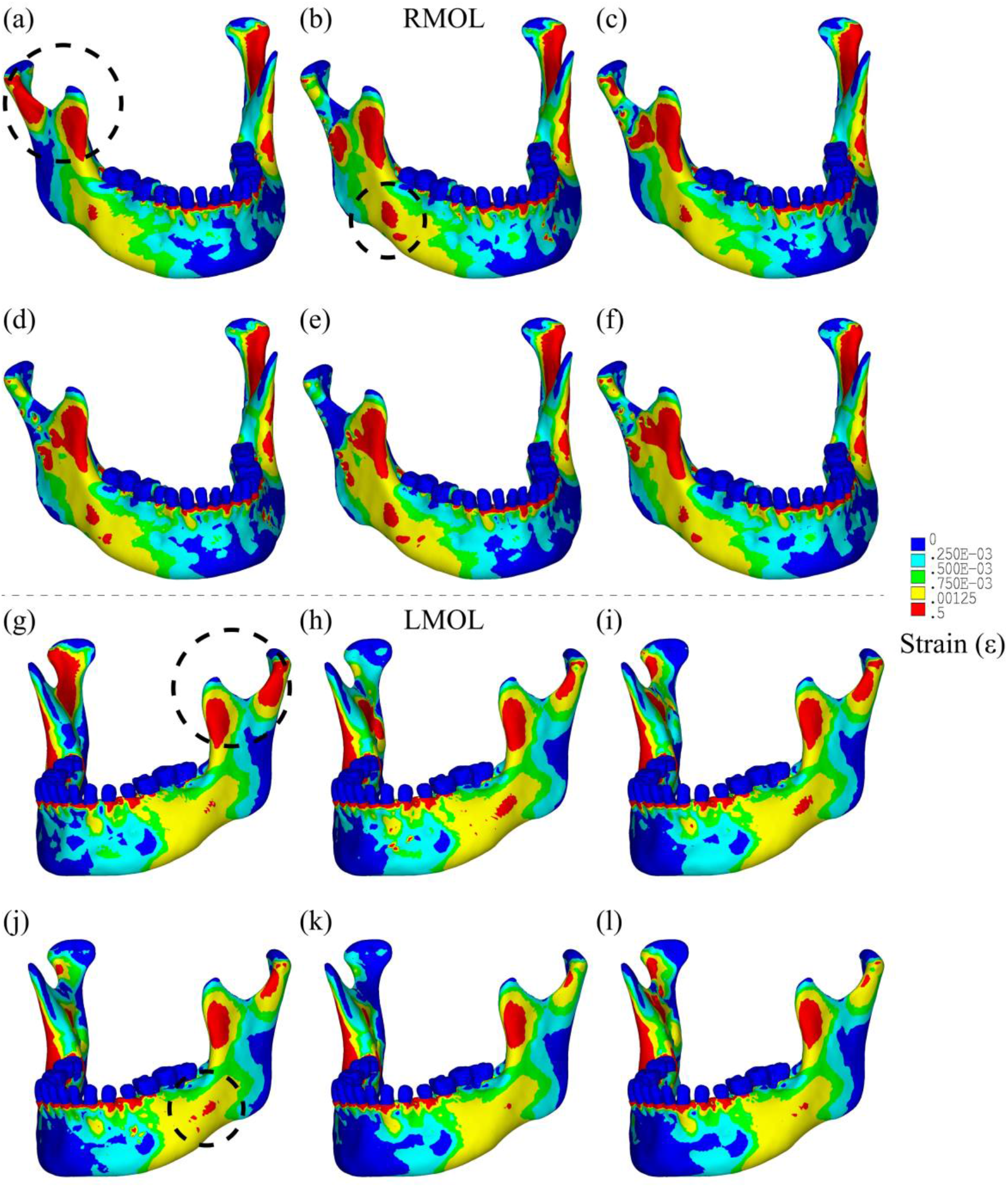
Principal tensile strain distributions: (a, g) intact and reconstructed mandibles with, (b, h) single mini, (c, i) trapezoid, (d, j) strut, (e, k) lambda, (f, l) double mini plates under RMOL and LMOL

As compared to the intact mandible, higher strains were observed in all the reconstructed mandibles during the molar clenching, along with considerable variation in the overall strain distribution with the clenching side (ipsilateral vs contralateral). Under ipsilateral occlusion (RMOL), the highest maximum tensile strain (1087 με) was observed in the reconstructed mandible with double mini plates. The strut plate caused the least maximum tensile strain (1039 με) in the mandible (Figure 5d). High compressive strains (920 – 970 με) were observed around the screw holes and near the anterior border of the mandibular coronoid (Figure S1b-f). The single mini plate resulted in the highest maximum compressive strains (~970 με) in the mandible (Figure S1b). The contralateral occlusion (LMOL) resulted in higher tensile strains in the reconstructed mandibles for all plates except the strut plate. Under LMOL, the mandible reconstructed with double mini plates (Figure 5l) experienced lower maximum tensile strain (1044 με) as compared to the reconstructed mandible with the single mini plate with the highest maximum tensile strain (1055 με) (Figure 5g). The lambda plate caused the least maximum tensile strain (1013 με) in the reconstructed mandible (Figure 5k). The trapezoid plate (Figure S2c) resulted in the least maximum compressive strain (910 με) in the reconstructed mandible.

### 3.2. Stresses in implants

For all the load cases, double mini plates exhibited the lowest von Mises stresses among all plates. Under RMOL among all plates, the highest maximum von Mises stress was found in the lambda plate (374 MPa) with high stresses below the screws at the central region (Figure 6). In comparison, there was a reduction in the maximum von Mises stresses in lambda plate under LMOL (316 MPa). On the contrary, the other plates exhibited a reverse trend, wherein, higher maximum von Mises stresses were observed during LMOL (~300 – 422 MPa) as compared to those during RMOL (170 – 255 MPa). Under LMOL, high stresses were observed in the anterior arm of the trapezoid and strut plates, and around the central screws in single and double mini plates (Figure 6).

**Figure 6.**
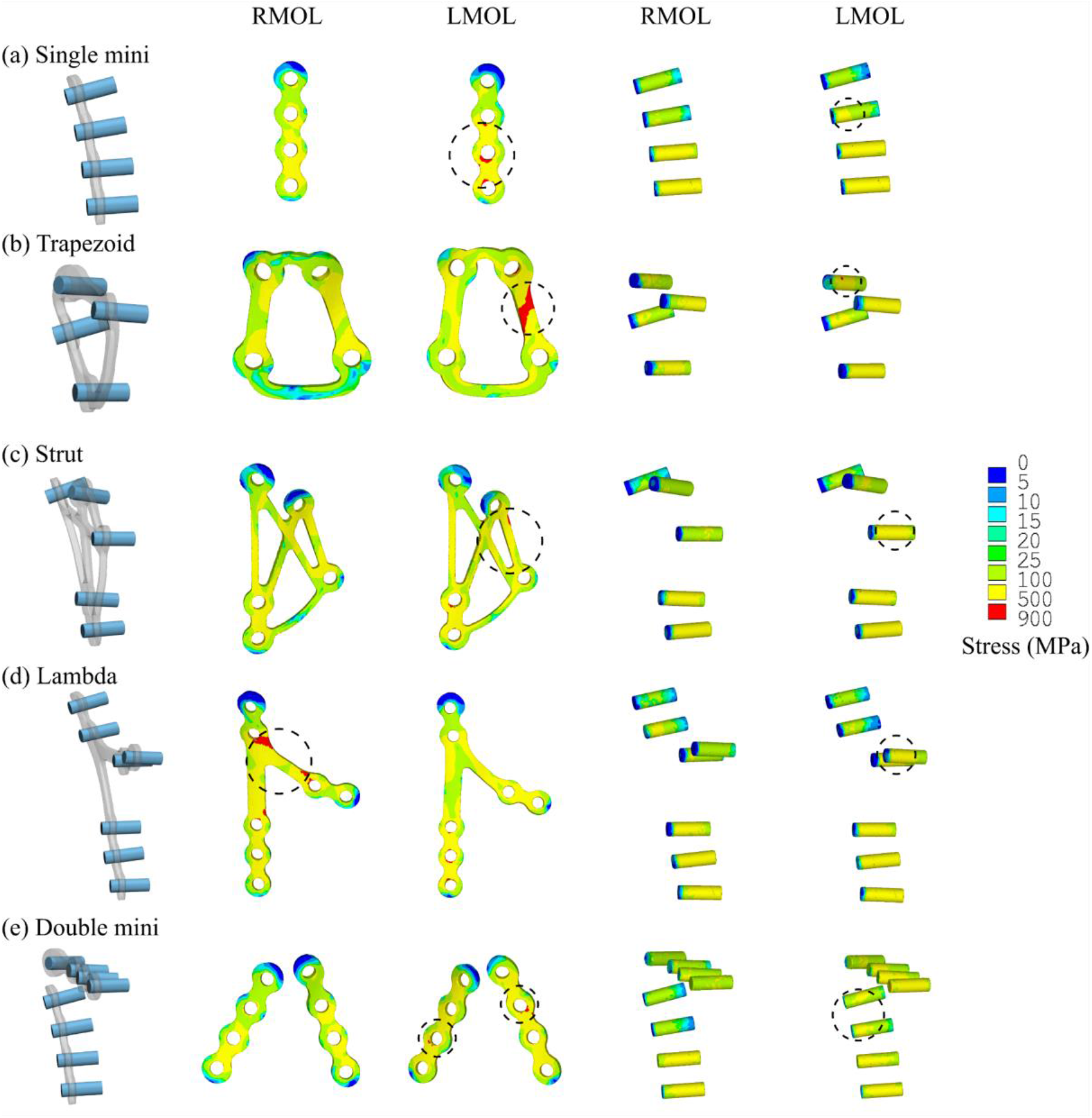
von Mises stress distributions: (a) single mini (b) trapezoid (c) strut (d) lambda (e) double mini plates under RMOL and LMOL

The maximum von Mises stresses in the screws were higher under LMOL (180 MPa – 300 MPa) than RMOL (130 MPa – 200 MPa) for all the plating techniques (Figure 9). Under LMOL, the maximum von Mises stresses in the plates were found to be inversely related to the number of screw holes (Figure 7). Under RMOL, the strut and lambda plate had higher von Mises stresses than the trapezoid plate. Maximum von Mises stresses in all the plates and screws during all clenching tasks was found to be lower than the yield strength of titanium alloy (~900 MPa^36^).

**Figure 7.**
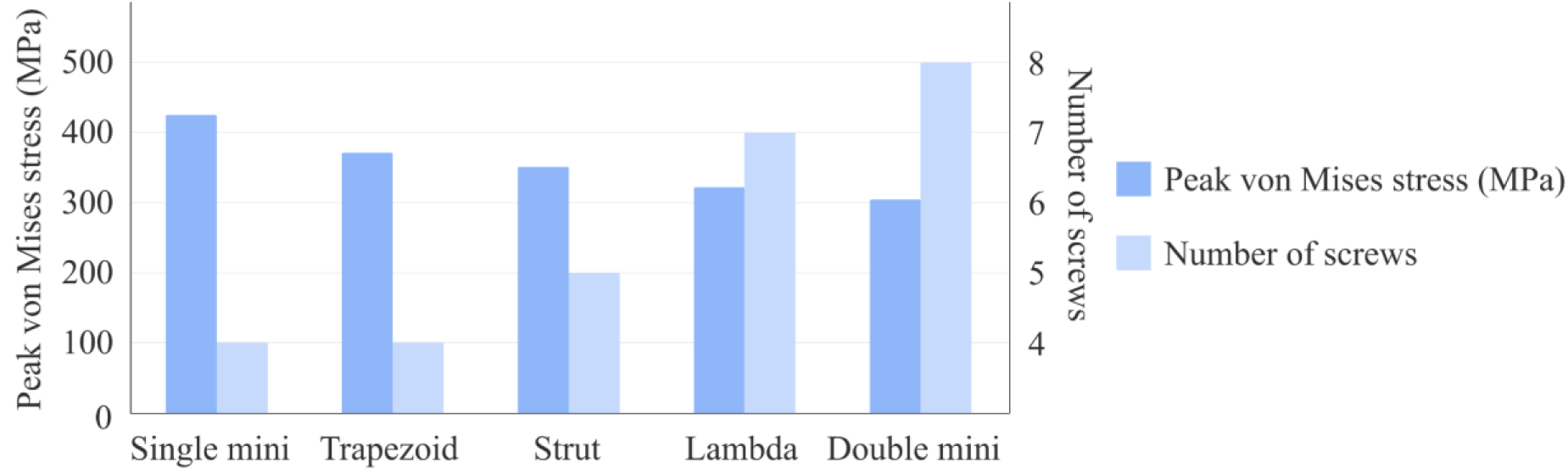
Comparison of von Mises stress in different plates under LMOL

### 3.3 Interfragmentary displacement

It was found that reconstruction with double mini plates led to the least interfragmentary gap after all the application of all the load cases. Under LMOL, the highest interfragmentary displacement of 200 μm was observed for the single mini plate. And under RMOL, the lambda plate resulted in the highest interfragmentary displacement of 300μm. A greater interfragmentary displacement was observed under LMOL as compared to RMOL, for all plates except the lambda plate, as described in Table 2.

**Table 2.**
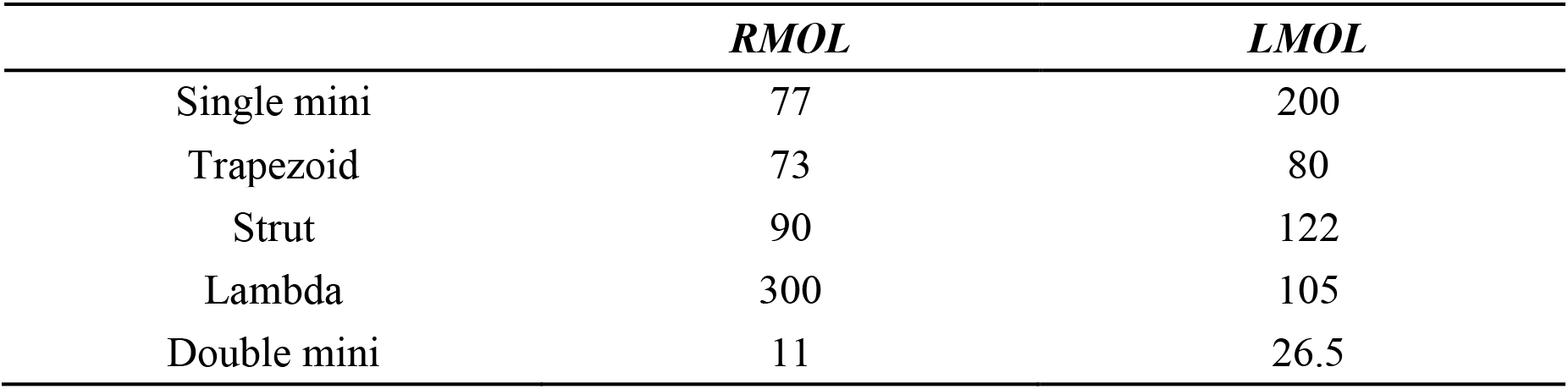
Interfragmentary gap (μm) during RMOL and LMOL

## 4. Discussion

Aimed at the identification of the most suitable design in terms of superior biomechanical fixation, the study sought to analyse the load transfer across intact and reconstructed mandibles with five different plate designs during a complete mastication cycle. The predictions of the developed FE model were qualitatively compared with previously published clinical and experimental studies. Despite the differences in the absolute magnitudes of stresses and strains ^21, 24^, there was overall similarity in terms of magnitudes and location of the strains and stresses in the mandible between our FE predictions and previous publications^21, 24^ which provide confidence in our FE predictions. The difference in the absolute magnitudes of stresses and strains between our FE predictions and previous publications could be attributed towards the variations in mandibular shape and material properties.

It was observed that the single mini plate resulted in a higher maximum tensile strain (1055 με) as compared to double mini (1044 με), under LMOL in the reconstructed mandible. This could possibly be due to the lack of neutralisation of tensile strain at anterior border^12^ with the single mini plate. However, under the ipsilateral clenching condition (RMOL), a reverse trend was observed, wherein, the single mini plate resulted in lower maximum tensile strain (1076 με) as compared to those (1087 με) of double mini plates (Figure 5b, f). Such a shift in the tensile strain contours with the clenching sides was also reported earlier^37^. The lowest compressive strains were observed with the trapezoid and the strut plates under LMOL and RMOL. This could be attributed to the fact that both trapezoid and the strut plates have large surface area and bridges between the anterior and posterior arms (Figure 2).

Among all the plate designs, the strut plate was found to be optimal during RMOL as the least amount of stresses and strains were observed in the reconstructed mandible. This might be due to the presence of anterior and posterior arms placed over tensile and compressive contours as well as the structural design with cross-bars (bridge) between the arms. These observations were similar to those reported by Murakami et al.^13^ wherein, the authors have reported the strut plate to be the most suitable configuration for unilateral condylar fracture.

The contralateral occlusion (LMOL) resulted in the higher peak von Mises (equivalent) stresses in plates and screws (~250-400 MPa vs ~150-200 Mpa) and lower strain in mandible (1013-1055 με vs. 1039-1087 με) as compared to ipsilateral molar clenching. Hence, the contralateral occlusion seems to be more suitable for the patients during immediate post-operative period and is also suggested clinically^38^. However, under the contralateral occlusion, the load transfer through the plates and screws increased due to the increase in the out-of-plane bending moment. Consequently, under the contralateral loading condition (LMOL), the single mini resulted in the highest interfragmentary displacement of 200 μm, which exceeded the limit of interfragmentary displacement (150 μm) considered ideal for fracture healing^39, 40^. On the contrary, the double mini and trapezoid plates exhibited notably reduced strains (~1043 με) and interfragmentary displacements of 26.5 μm and 80 μm, respectively.

In addition to the single mini plate resulting in the highest interfragmentary displacement among all the plates, it also had higher peak von Mises stress as compared to double mini plate in all the load cases. This evaluation also conforms to the literature reports that single mini plate is not favourable^7–9, 13^. However, all the plates could safely sustain the loads during the mastication cycle, as the peak von Mises stresses were within the yield limit, thus Ti-6Al-4V suffices the fixation requirements of sustaining the physiological loads.

However, there were certain limitations of the present study. Although both cancellous and cortical exhibit heterogeneous and anisotropic material behaviour, the heterogeneous isotropic material properties were assigned to the cancellous and region-wise homogeneous orthotropic material properties were assigned to the cortical. Another limitation was that the subcondylar fracture was created without gap for ideal clinical conditions representing a complete open-reduction internal fixation procedure. Inter-patient variability in the muscle forces during the mastication cycle was also not considered here. Therefore, the results of this comparative study are patient-specific and should be interpreted within the limitations.

In summary, the study examined the influence of different plate configurations on load transfer through mandible during different clenching tasks. The contralateral molar clenching task is preferred during the immediate post-surgery period for patients. Under this contralateral molar clenching, the peak von Mises stresses in the plate decreased with increase in the number of screws. Furthermore, the presence of two arms seems beneficial to neutralise the tensile and compressive strains across load cases. Consequently, double mini and trapezoid plates were found to perform better as compared to single mini plate during the entire mastication cycle for subcondylar fracture fixation.

