## supplementary information for "Biomechanical influence of plate configurations on mandible subcondylar fracture fixation: a finite element study"

**SUPPLEMENTARY MATERIAL**

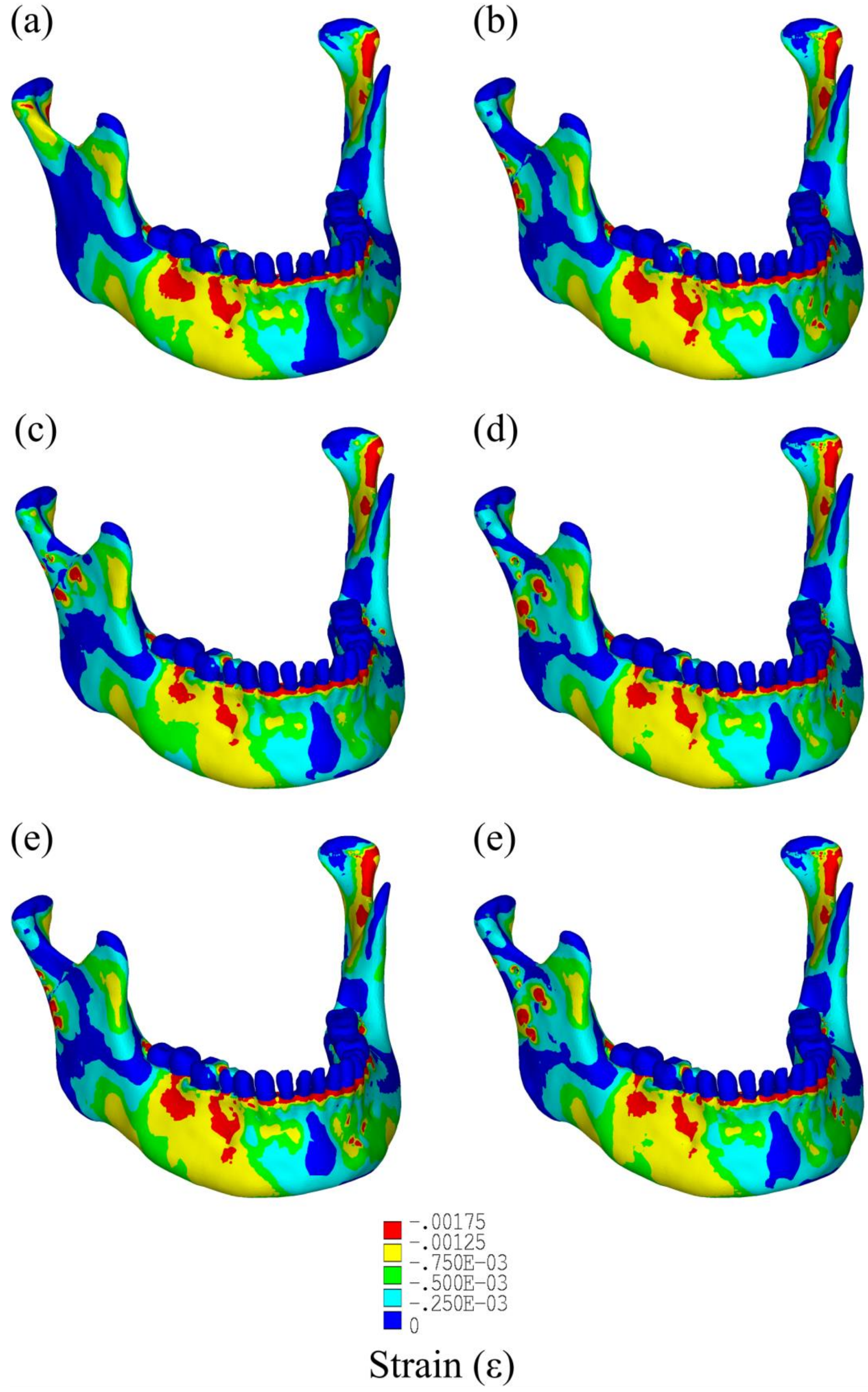


**Figure S1.** Principal compressive strain distributions: (a) intact and reconstructed mandibles with, (b) single mini (c) trapezoid (d) strut (e) lambda (f) double mini plates under RMOL

### SUPPLEMENTARY MATERIAL

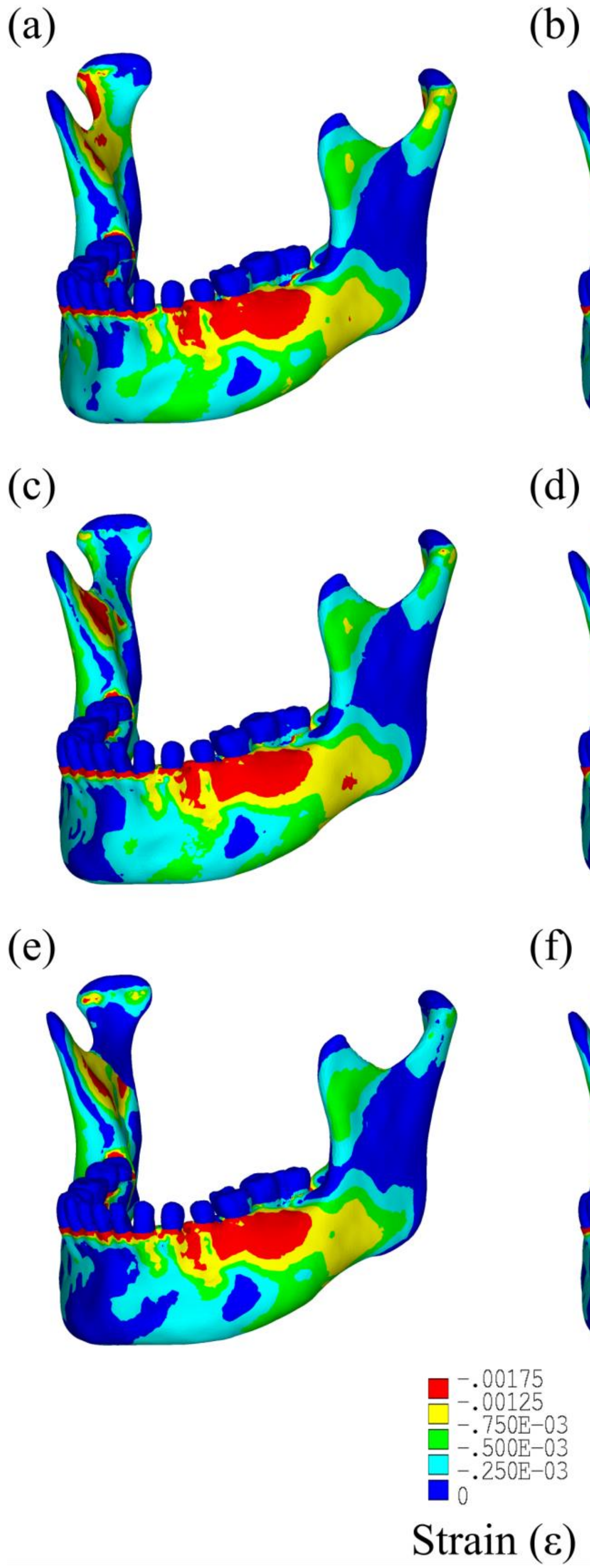


**Figure S2.** Principal compressive strain distributions: (a) intact and reconstructed mandibles with, (b) single mini (c) trapezoid (d) strut (e) lambda (f) double mini plates under LMOL

### SUPPLEMENTARY MATERIAL

**Table S1.** Maximum tensile stress (MPa) for intact and reconstructed mandibles

| *Load case* | *RMOL* | *LMOL* | *RGF* | *LGF* | *INC* | *ICP* |
|---|---|---|---|---|---|---|
| Healthy | 14.77 | 14.13 | 11.52 | 7.94 | 9.38 | 15.87 |
| Trapezoid | 18.12 | 15.89 | 10.26 | 9.04 | 11.53 | 17.16 |
| Single mini | 17.41 | 15.79 | 9.815 | 8.9 | 11.46 | 17.31 |
| Strut | 17.14 | 15.04 | 9.78 | 8.79 | 11.09 | 16.84 |
| Lambda | 16.94 | 15.16 | 9.405 | 8.67 | 10.92 | 16.87 |
| Double mini | 17.29 | 15.45 | 9.73 | 8.74 | 11.14 | 16.65 |

**Table S2.** Maximum tensile strain (με) for intact and reconstructed mandibles

| *Load case* | *RMOL* | *LMOL* | *RGF* | *LGF* | *INC* | *ICP* |
|---|---|---|---|---|---|---|
| Healthy | 940 | 947 | 495 | 478 | 514 | 988 |
| Trapezoid | 1069 | 1042 | 568 | 536 | 652 | 1124 |
| Single mini | 1076 | 1055 | 581 | 581 | 622 | 1182 |
| Strut | 1039 | 1054 | 577 | 543 | 625 | 1118 |
| Lambda | 1054 | 1013 | 560 | 520 | 637 | 1087 |
| Double mini | 1087 | 1044 | 564 | 550 | 665 | 1119 |

**Table S3.** Maximum compressive strain (με) for intact and reconstructed mandibles

| *Load case* | *RMOL* | *LMOL* | *RGF* | *LGF* | *INC* | *ICP* |
|---|---|---|---|---|---|---|
| Healthy | 864 | 870 | 448 | 441 | 472 | 780 |
| Trapezoid | 940 | 910 | 506 | 532 | 577 | 918 |
| Single mini | 969 | 958 | 541 | 493 | 557 | 976 |
| Strut | 918 | 913 | 515 | 496 | 534 | 894 |
| Lambda | 920 | 927 | 481 | 495 | 571 | 892 |
| Double mini | 948 | 962 | 488 | 516 | 575 | 894 |

### SUPPLEMENTARY MATERIAL

**Table S4.** Maximum von Mises stress (MPa) in the plates

| *Load case* | *RMOL* | *LMOL* | *RGF* | *LGF* | *INC* | *ICP* |
|---|---|---|---|---|---|---|
| Trapezoid | 213.6 | 382.2 | 218.54 | 220.4 | 235.66 | 328.59 |
| Single mini | 244.3 | 422.85 | 201.19 | 219.98 | 231.34 | 289.14 |
| Strut | 254.9 | 358.37 | 363.47 | 218.74 | 202.41 | 275.29 |
| Lambda | 374 | 315.83 | 164.88 | 248.05 | 268.17 | 267.28 |
| Double mini | 178.1 | 302.22 | 123.99 | 172.57 | 178.67 | 242.41 |

**Table S5.** Maximum von Mises stress (MPa) in the screws

| *Load case* | *RMOL* | *LMOL* | *RGF* | *LGF* | *INC* | *ICP* |
|---|---|---|---|---|---|---|
| Trapezoid | 148.5 | 298.23 | 109.36 | 162.17 | 161.24 | 201.69 |
| Single mini | 199.3 | 271 | 104.84 | 139.57 | 153.008 | 171.37 |
| Strut | 168.7 | 238.3 | 107.35 | 139.3 | 150.72 | 175.28 |
| Lambda | 173.4 | 237 | 128.42 | 159.64 | 174.66 | 182.19 |
| Double mini | 132.3 | 180.8 | 86.4 | 104.05 | 109.92 | 131 |

**Table S6.** Interfragmentary displacements (μm)

| *Load case* | *RMOL* | *LMOL* | *RGF* | *LGF* | *INC* | *ICP* |
|---|---|---|---|---|---|---|
| Trapezoid | 73 | 80 | 100 | 37 | 44 | 104 |
| Single mini | 77 | 200 | 120 | 36 | 40 | 140 |
| Strut | 90 | 122 | 340 | 140 | 60 | 160 |
| Lambda | 300 | 105 | 55 | 106 | 120 | 197 |
| Double mini | 11 | 26.5 | 11.8 | 9 | 10 | 27 |
